# A comparative microarray analysis identifies a conserved gene expression signature between airway injury and lung cancer

**DOI:** 10.1101/090605

**Authors:** Adam Giangreco

## Abstract

Lung squamous cell carcinoma (SqCC) accounts for 30% of lung cancers, with over 400,000 deaths per year worldwide. Although evidence suggests that chronic lung injury drives carcinogenesis, a comprehensive understanding of this process remains elusive. Here, I used a comparative microarray analysis to identify gene expression differences shared between airway injury and squamous lung cancer. Of the 667 genes that exhibited differential expression following murine polidocanol and SO2 injury, 40.6% were additionally dysregulated in human SqCC. Among these, 150 genes were consistently upregulated and 54 downregulated relative to all controls. Examples included genes associated with increased cell cycling, aberrant cytokinesis and DNA repair, and enhanced tumour cell invasion and metastases. For 88.2% of identified genes, altered expression was associated with increased SqCC progression and patient mortality. These results establish a novel gene expression signature linking airway injury and lung cancer pathogenesis.

## Introduction

Lung cancers are the leading cause of cancer death in the world, with squamous cell carcinoma (SqCC) exhibiting a mere 14% 5-year survival while accounting for > 30% of lung cancer mortality ^1^. Lung SqCC occurs almost exclusively in smokers, is characterised by a very high mutational burden, and expresses abundant cytokeratins and squamous differentiation markers including cytokeratin 5 and TP63 ^2^. SqCCs often derive centrally from large airways, and in their early stages exhibit histological features reminiscent of chronic lung injury including basal cell hyperplasia ^3^. In contrast to lung adenocarcinomas, current treatment options for SqCC remain limited, with targeted chemotherapies against EGFR, HER2, and ALK proving largely ineffective ^4,5^. In addition, there is a paucity of clinically relevant biomarkers for predicting overall patient mortality, disease progression, and therapeutic response ^6^.

In recent years, high throughput sequencing has raised the possibility of cataloguing all mutations and chromosomal aberrations present within human lung SqCC ^7^. The aim of this research has been to identify mutations suitable for cancer risk prediction, to facilitate early clinical diagnosis, and to develop novel targeted therapies. Although a plethora of candidate tumour suppressors and oncogenes for SqCC have recently emerged, the importance of these for disease progression remains elusive. Contributing to this are the observations that TP53 remains the only identified oncogenic mutation present in the majority of SqCC patients ^8^, that only a subset of early SqCC lesions progress to invasive disease ^9,10^, and that significant intra-tumoral genetic heterogeneity is present in most disease^11^. Thus, an increased understanding of the molecular alterations driving early lung carcinogenesis is needed.

Recently, genetically engineered mouse models have demonstrated that endogenous airway stem cells function as the cell-of-origin for murine lung cancer ^3,12,13^. In these models, it is notable that the incidence, speed, and severity of lung tumours are all increased following lung injury and the presence of a pro-inflammatory microenvironment ^14,15^. These results suggest that conserved, endogenous signalling pathways responsible for lung injury outcomes may additionally regulate lung tumorigenesis. To test the hypothesis that conserved changes in gene expression accompany both lung injury and cancer I performed a comparative microarray analysis of murine lung injury and human SqCC. This approach identified 204 genes whose expression was consistently altered in the presence of both murine lung injury and human SqCC. These 150 consistently upregulated and 54 downregulated genes included established regulators of cell cycling, DNA repair, and cell invasion, and were associated with increased SqCC progression and patient mortality.

## Results

### Murine polidocanol and SO2 injury models elicit conserved changes in gene expression

An overview of the comparative microarray analysis strategy is illustrated in Figure 1. To elucidate whether distinct murine lung injury models invoke conserved gene expression signatures, the profiles of murine tracheal samples isolated 48 hours post-polidocanol or post-SO2 injury were compared ^16,17^. Comparing these two different models facilitates the exclusion of genes whose differential expression is dependent solely on the type of injury. For each microarray (GSE17268, GSE69056) all probes with an adjusted p-value <0.05 and logFC between -0.5 and 0.5 were excluded, resulting in elimination of 70-95% of all genes. Conserved genes with a p<0.05 and logFC <-0.5 or >0.5 were identified using a ‘Find Matches’ Visual Basic macro, with those present within each microarray classified as ‘shared’. All shared genes exhibiting consistent up- or down-regulation were classified as ‘coregulated’ (see Supplemental Methods). Although this approach utilises previously published gene expression datasets, the current analysis techniques differ from those used in each previous study, resulting in significant differences in identified genes. An identical strategy for identifying differentially expressed and consistently up-and down-regulated genes was used to compare murine lung injury and human SqCC microarrays. An exact statistical test was used to compare the expected degree of overlap and intersection amongst each of these independent microarray datasets ^18^. This analysis predicted the number of genes that were likely to be present in comparative microarray dataset lists by chance alone (see Supplemental data). Using this approach, all identified gene lists were highly statistically significant (p<0.000001).

**Figure 1.**
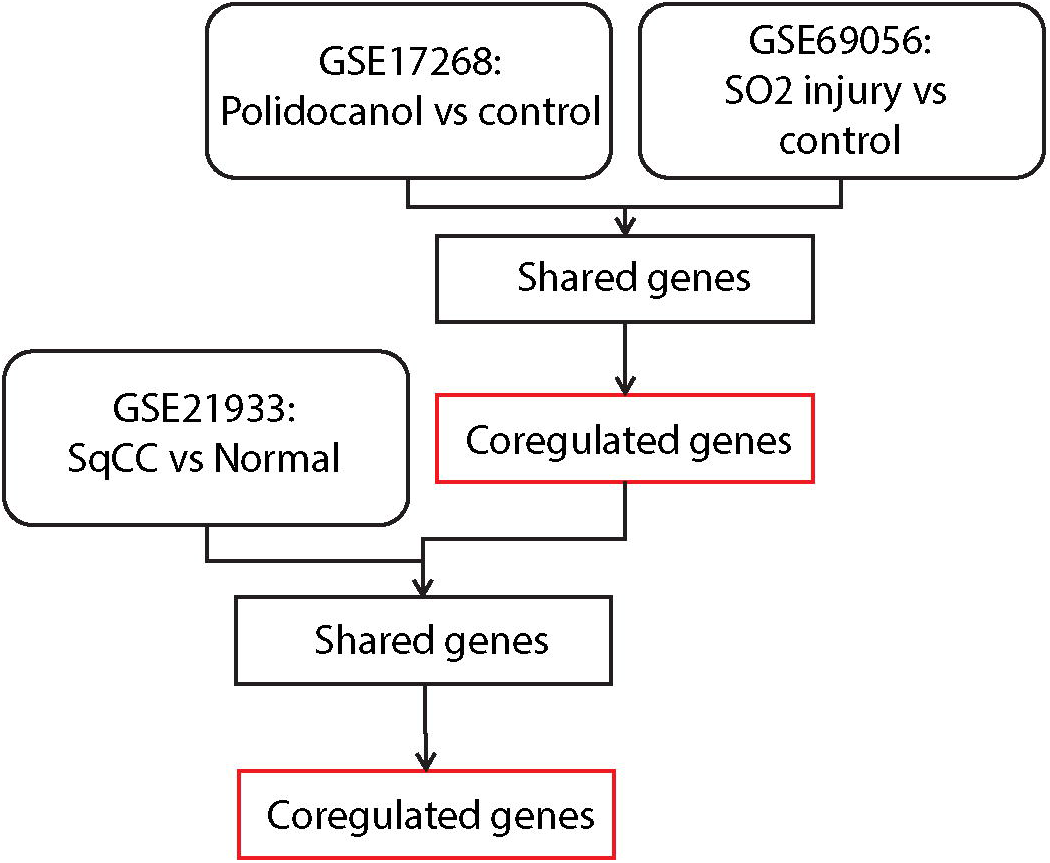
Flowchart for comparative microarray analysis. The identification of differentially expressed genes was assessed individually for each microarray platform (GSE17268, GSE69056, GSE21933). A comparative microarray analysis of probes exhibiting a p-value of <0.05 and logFC <-0.5 or >0.5 was performed to identify conserved differentially expressed genes (classified as ‘shared’). Shared genes exhibiting consistent up- or down-regulation relative to controls were further defined as ‘coregulated’. A statistical test for the significance of the overlap in genes identified in each of these independent microarray dataset comparisons is included in supplemental data.

Polidocanol microarrays involved comparison of 5 uninjured and 5 polidocanol treated mice and revealed 2222 differentially expressed probes (blue, Figure 2A, ^17^). SO2 microarray comparisons of 3 control and 3 SO2 injured animals revealed 8872 differentially expressed probes (yellow, Figure 2A, ^16^). The greater number of differentially expressed probes following SO2 injury (8872) may reflect the limited sample size of this analysis (n=3, ^16^). 1108 probes (representing 717 unique genes) were differentially expressed following both polidocanol and SO2 injury (green, Figure 2A). LogFC expression analysis indicated that 385 of these 717 genes (53.6% of total) were consistently upregulated in both polidocanol and SO2 injury while 282 genes (39.3% of total) were consistently downregulated (Figure 2B). Overall, these 667 consistently up and downregulated genes represent 93% of all 717 differentially expressed genes, confirming that murine polidocanol and SO2 injury invoke conserved changes in gene expression. The number of genes predicted to be differentially expressed in both microarray datasets by chance alone was 162, making these results extremely statistically significant (see Supplemental Data). Consistently upregulated genes were associated with increased proliferation and included Cyclins B1, B2, and A2 (Ccnb1, Ccnb2, Ccna2), cyclin dependent kinase 1 (Cdk1), Centrosomal protein 55kD (Cep55), polo-like kinase 1 (Plk1), and pro-platelet basic protein (Ppbp/Cxcl7, Figure 2C and Supplemental Figure 1. Downregulated genes included Flavin monooxygenase 3 (Fmo3), paraoxonase 1 (Pon1), aldehyde oxidase 3 (Aox3), and Cell adhesion molecule 1 (Cadm1, Figure 2D and Supplemental Figure 2). A complete list of all 667 consistently up and downregulated genes and their logFC expression values are included in Supplemental Table 2.

**Figure 2.**
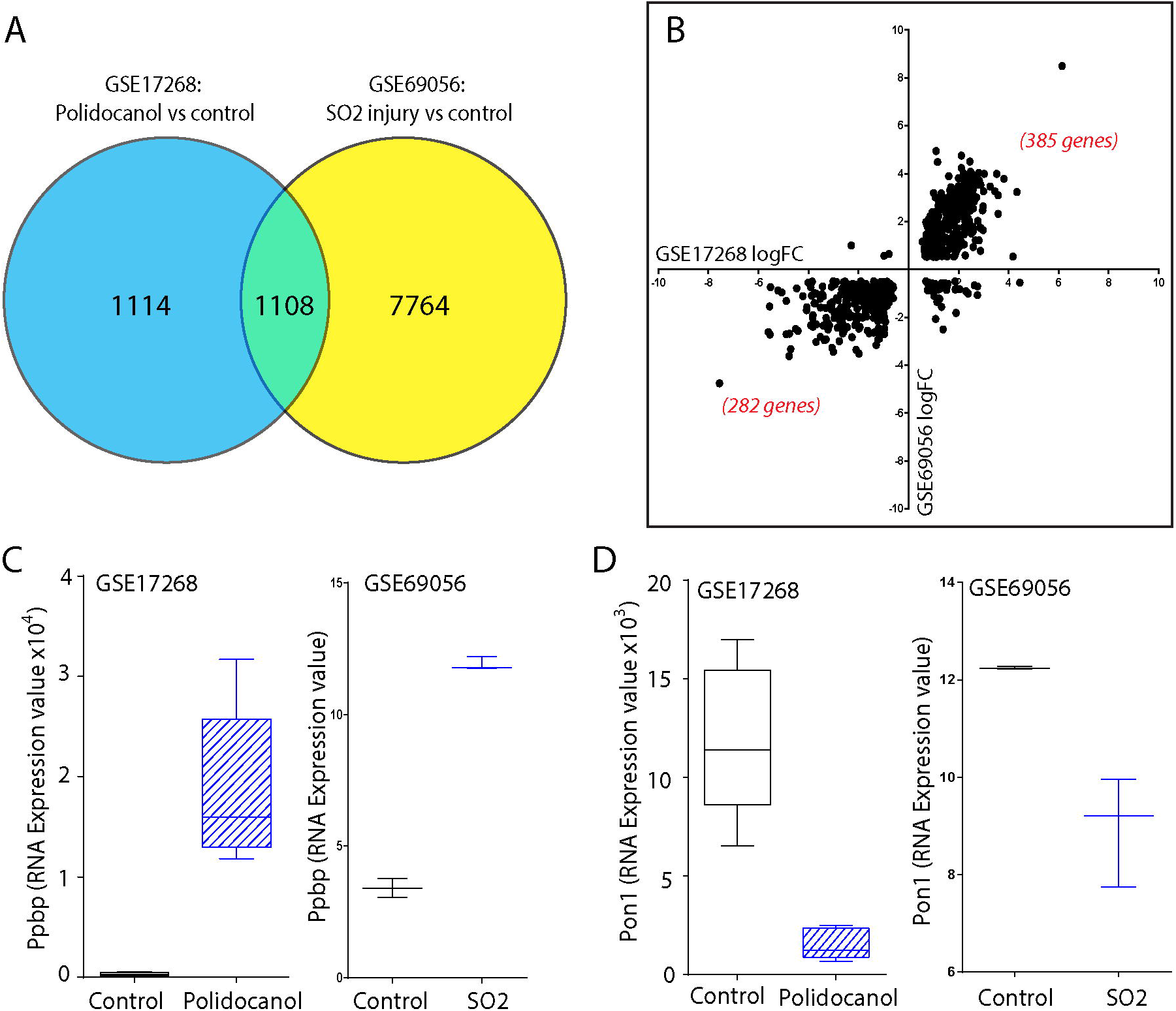
Comparative microarray analysis identifies a shared BSC-mediated repair signature following polidocanol and SO2 exposure. (A) Venn diagram of comparative microarray analysis between polidocanol (GSE17268) and SO2-injured airways (GSE69056). Green denotes 1108 probes representing 717 individual genes exhibiting differential expression in both microarray datasets (logFC <-0.5 or >0.5; p<0.05). Blue and yellow identify differentially expressed probes exclusive to GSE17268 or GSE69056. (B) Analysis of logFC expression values for all 717 conserved differentially expressed genes present in both datasets. (C, D) Individual examples of consistently upregulated and downregulated genes following polidocanol and SO2 injury include pro-platelet basic protein (Ppbp/Cxcl7, C) and paraoxonase 1 (Pon1, D). A complete list of differentially expressed genes and their logFC expression values is provided in Supplemental Table 2.

### Comparative microarrays identify a conserved gene expression signature between airway injury and squamous carcinogenesis

To determine whether changes in gene expression following murine lung injury also exhibit differential expression in human lung cancer I compared all 667 differentially expressed genes against 8271 differentially expressed human SqCC probes (GSE21933, Figure 3A). Results indicate that 271 genes were differentially expressed in both murine injury and human SqCC, representing 40.6% of all 667 consistently up and downregulated genes in murine tracheal injury (green, Figure 3A). Of these 271 genes, 168 were consistently upregulated and 66 downregulated, representing 61.9% and 24.3% of all conserved differentially expressed genes in murine tracheal injury and human SqCC, respectively (Figure 3B). The differential expression of this 234 gene subset was further validated using an independent cohort of 65 normal and 27 SqCC tumour biopsy samples (GSE19188, ^19^). Overall, 89.2% (150 out of 168) of upregulated genes and 81.8% (54/66) of downregulated genes were validated using this independent cohort. These 204 genes represent 30.5% of all 667 consistently up and downregulated differentially expressed genes identified following murine lung injury, demonstrating the high degree of conservation in gene expression between injury and carcinogenesis. The number of genes predicted to be differentially expressed in all four microarray datasets by chance alone was 0.897, making these results extremely statistically significant (see Supplemental Data).

**Figure 3.**
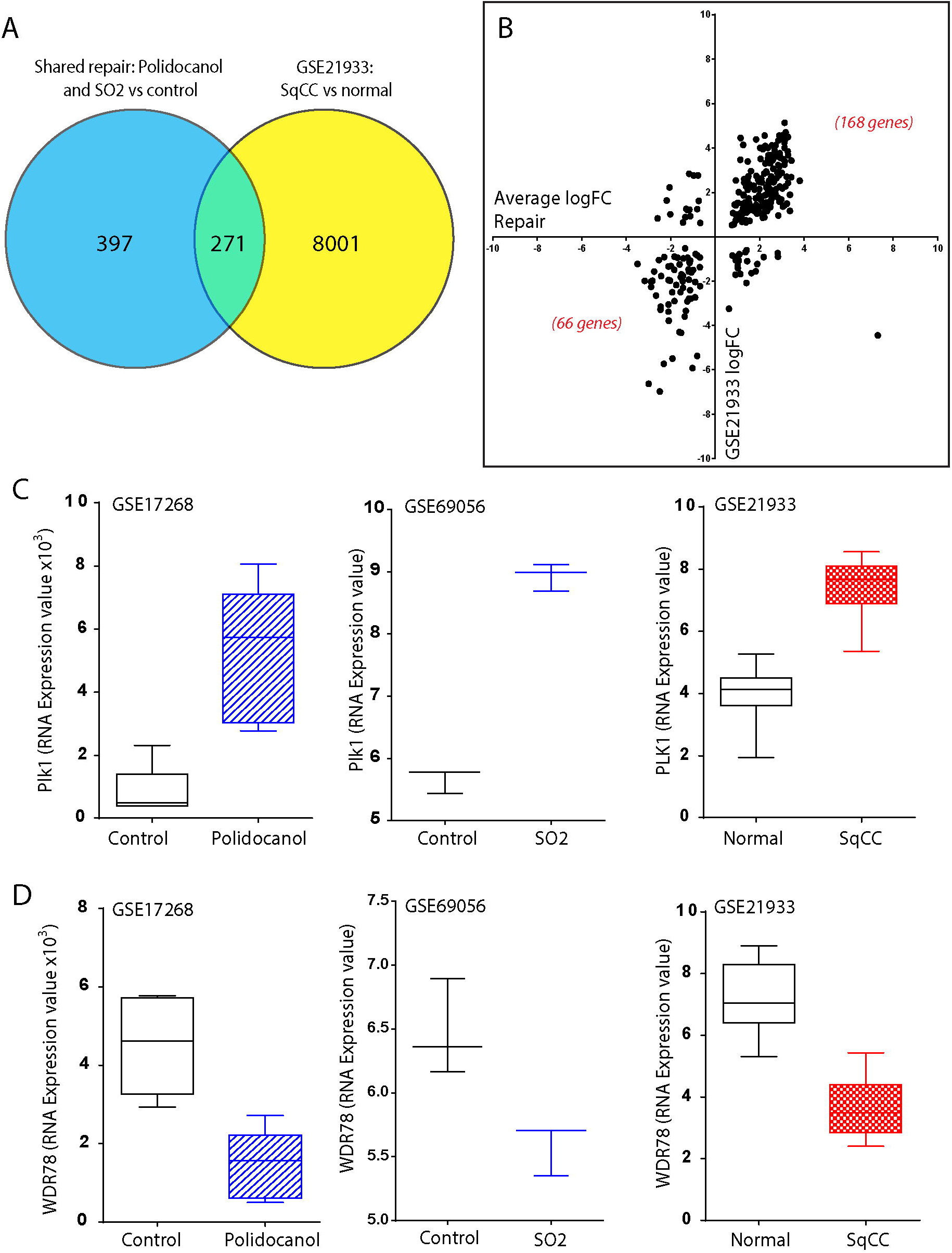
Identification of repair signature genes conserved in human lung SqCC. (A) Venn diagram of comparative microarray analysis between shared, coregulated genes of airway repair (blue) and genes differentially expressed in human SqCC (yellow, GSE21933). Green denotes 271 individual genes exhibiting differential expression in both datasets (logFC <-0.5 or >0.5; p<0.05). (B) Analysis of the log FC expression values for all 271 differentially expressed genes present in airway repair and human SqCC, of which 204 were further validated using an independent human SqCC gene microarray dataset (GSE19188). (C, D) Examples of these 204 consistently upregulated and downregulated genes in airway repair and SqCC include polo-like kinase 1 (PLK1, C) and WD-containing repeat domain 78 (WDR78, D). A complete list of all these 204 independently validated, coregulated genes and their logFC expression values is provided in Supplemental Table 3. Additional examples of individual up- and down-regulated genes are provided in Supplemental Figures 1 and 2.

Examples of upregulated genes include the transcription factor FOXM1, its target genes centrosomal protein 55 (CEP55) and helicase lymphoid-specific (HELLS), and PLK1 (Figure 3C and Supplemental Figure 1). Additional upregulated genes included ubiquitin-like with PHD and ring finger domains 1 (UHRF1), ribonuclease reductase subunit M2 (RRM2), deoxythymidylate kinase (DTYMK), RAD51, and RAD54 (Supplemental Figure 1). Consistently downregulated genes included WDR78 (Figure 3D), the known tumour suppressors Cell adhesion molecule 1 (CADM1, ^20^) and aldehyde dehydrogenase 2 (ALDH2, ^21^), and several genes related to oxidoreductase activity that have recently been described as pan-cancer biomarkers (including AOX1 and ACOX2, Supplemental figure 2, ^22^). In addition, biomarkers of lung epithelial differentiation including surfactant proteins B, C, and D (SFTPB, SFTPC, SFTPD), flavin monooxygenases 2 and 3 (FMO2, FMO3), and the lung-specific cytochrome p450 isotype B1 (CYP4B1) were downregulated in both lung injury and squamous lung cancer. A complete list of all 204 independently validated, consistently up- and down-regulated genes is included in Supplemental Table 3.

### Conserved gene expression changes predict SqCC patient mortality and disease progression

To examine whether identified changes in gene expression that are conserved between murine lung injury and human carcinogenesis correlate with human SqCC mortality, I compared patient survival against relative gene expression for all 204 consistently up and down regulated genes using a KMplot analysis tool ^23^. Kaplan-Meier analyses revealed that 88.2% (180/204) of identified gene expression changes were associated with significantly altered patient survival outcomes. As individual examples of highly differentially expressed genes associated with altered patient survival, PLK1 and WDR78 were selected on the basis of their robust, consistent up and downregulation across both murine tracheal injury and human SqCC datasets (see Supplemental Tables 2 and 3). As expected, SqCC patients with high PLK1 expression had elevated mortality relative to those with low PLK1 expression (p<1E-16, HR = 2.23; Figure 4A). Conversely, patients with low WDR78 expression exhibited reduced mortality, consistent with the downregulation of this gene in murine lung injury and human SqCC (p<5.9E-12, HR = 0.53; Figure 4B). In addition to highly differentially expressed genes such as PLK1 and WDR78, consistently up and down regulated genes with much less robust differential expression also exhibited significant differences in overall patient survival. Examples included the consistently upregulated gene POLA2 (p=3.8E-12, HR=1.56; Figure 4C) as well as the consistently downregulated gene RABGAP1L (p=5.6E-11, HR=0.62; Figure 4D). Overall, these results suggest that the vast majority (88.2%, 180/204) of consistently differentially expressed genes can serve as predictors of patient mortality, regardless of their degree of differential expression. These findings also raise the possibility that identified genes and their corresponding signalling pathways represent functionally significant targets for human SqCC progression.

**Figure 4.**
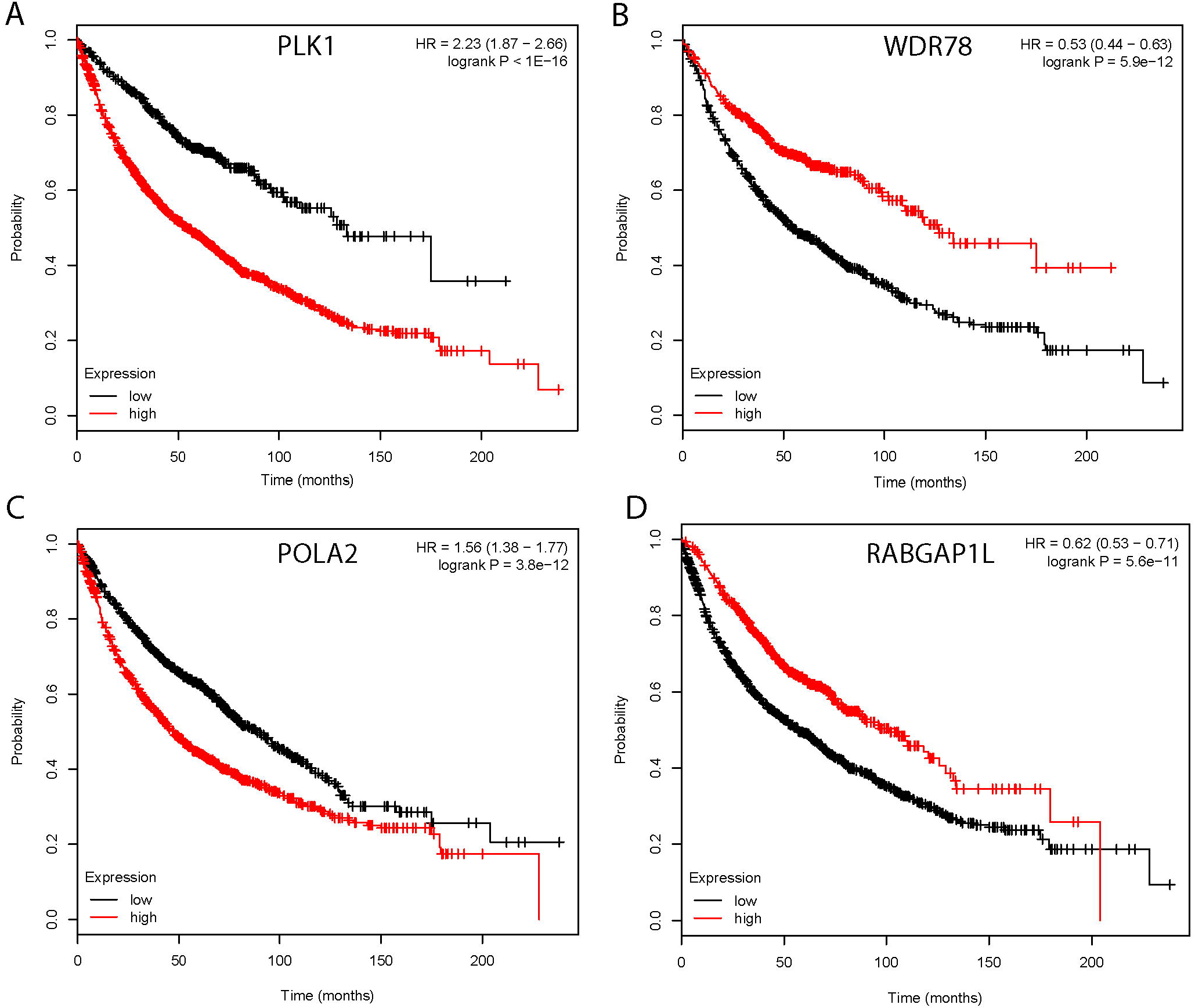
Conserved biomarker expression correlates with human NSCLC survival. Kaplan-Meier survival analysis was used to determine the relationship between all upregulated and downregulated gene expression levels with overall NSCLC patient survival. Overall, 88.2% (180/204) of all consistently differentially expressed genes were associated with significantly altered patient mortality. Individual examples of (A, C) upregulated (PLK1, POLA2) and (B, D) downregulated (WDR78, RABGAP1L) genes are shown along with their relative hazard ratio (HR) and logrank P value. Data was assessed using the online KMplot tool (www.kmplot.com).

To assess whether identified changes in gene expression may be associated with early human SqCC progression I examined one of the most significantly and consistently upregulated genes, PLK1, using a cohort of longitudinally collected bronchoscopic biopsy samples exhibiting various stages of preinvasive disease ^9^. PLK1 was selected for this additional analysis on the basis of its robust upregulation in both tracheal injury and human SqCC datasets (Supplemental Tables 2 and 3) as well as its strong correlation with increased patient mortality (Figure 4A). Samples for PLK1 immunostaining were obtained from patients under surveillance for lung cancer and represent a spectrum of normal, dysplastic, carcinoma-in-situ (CIS), and invasive SqCC phenotypes. In all patients, epithelial cells were identified using the pan-cytokeratin antibody AE1/3 (green, Figure 5), and preinvasive lesions underwent progression within 1 year. In normal biopsy samples, low amounts of PLK1 were detectable throughout all airway epithelial cells (red, Figure 5). In contrast, samples classified as preinvasive dysplasia and carcinoma in situ appeared to exhibit increased PLK1 staining intensity at levels similar to invasive SqCC (red, Figure 5). As a comparison, the abundance of AE1/3 immunostaining was consistent at each tumour stage (green, Figure 5). These results suggest that increases in PLK1 protein abundance, identified via indirect immunostaining, may represent an early event in SqCC pathogenesis.

**Figure 5.**
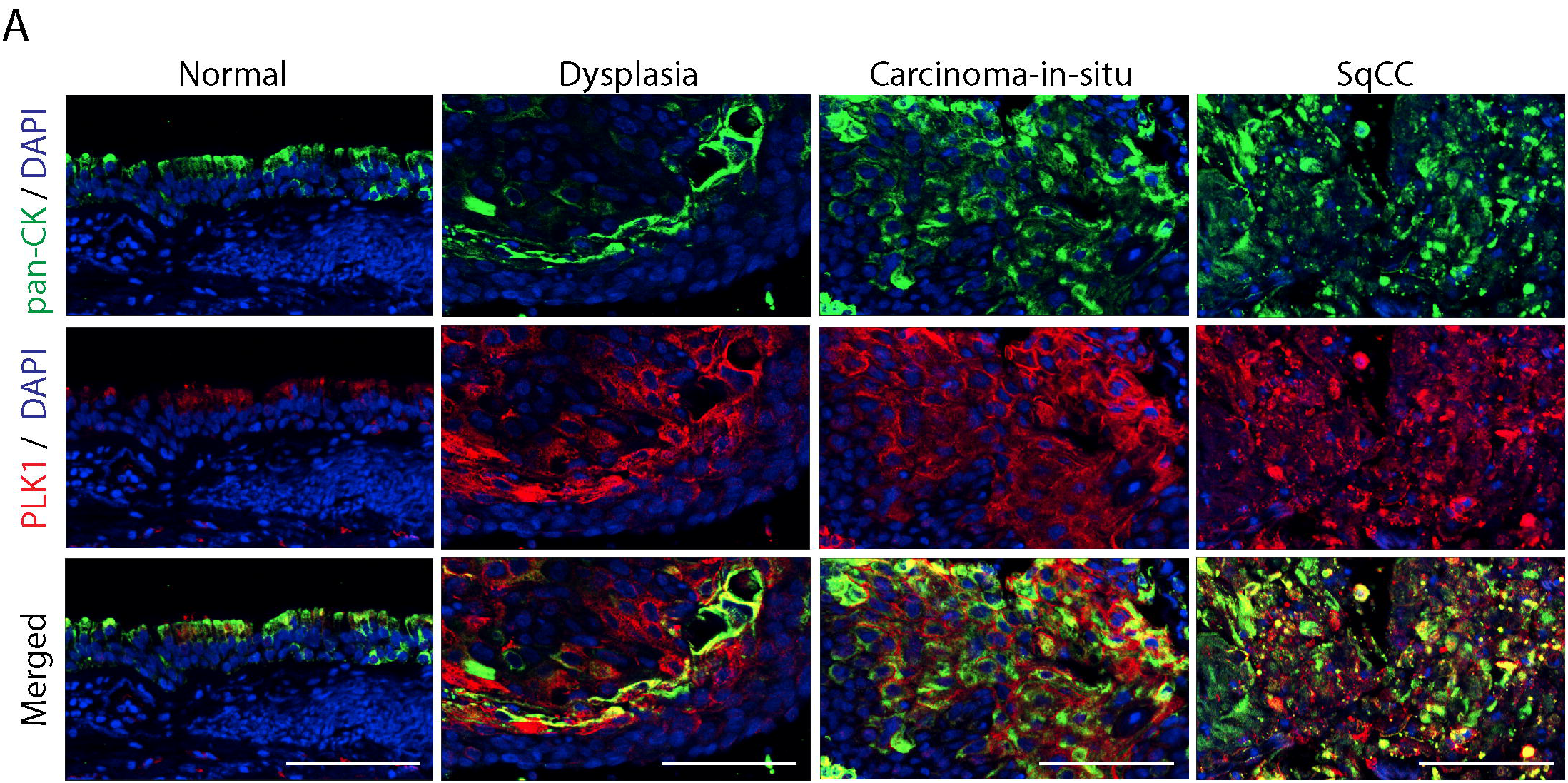
A conserved biomarker of airway repair and carcinogenesis is upregulated in preinvasive SqCC lesions. Immunofluorescent staining of PLK1 (red) plus pan cytokeratin (AE1/3, green) in normal, dysplastic, carcinoma-in-situ, and invasive SqCC biopsy samples. Staining shows increased expression of PLK1 in dysplastic, preinvasive, and SqCC samples compared with expression in the normal epithelium. DAPI is used as a nuclear counterstain (blue), and scale bars denote 100μm.

## Discussion

In this study, a combination of microarray analysis, immunostaining, tumour staging, and patient survival data identified a conserved gene expression signature linking airway injury and lung cancer. This included 150 upregulated and 54 downregulated genes that provide insights into the molecular alterations that link lung injury with lung tumour development. Consistently up- and down-regulated genes included those linked to transcriptional pathways necessary for cell cycling, cytokinesis, and DNA repair. Given the nature of the microarray experiments used in this study, there remain several potential explanations for the presence of these consistently differentially expressed genes. One possibility is that similarities in the stem cells responsible for both lung repair and tumourigenesis drive similar gene expression changes, as has been recently described using various mouse tumour models^2,24,25^. Additonal mechanisms to explain this overlap include the possibility that injury and cancer induce similar changes to the surrounding microenvironment, including differences in stromal, immune, and inflammatory cell phenotype and abundance as well as altered extracellular matrix stiffness and composition. Regardless of the origin of these changes, the discovery that 88.2% (180/204) of them are associated with altered lung cancer patient survival demonstrates the clinical significance of this approach, and suggests that a high percentage of differentially expressed genes may also be functionally relevant. It may also be possible to use consistently differentially expressed genes for predicting patient survival and risk of disease progression.

A significant number of the genes identified in this study are known regulators of a variety of human cancers, and several of these have now begun to be investigated therapeutically. Of note, targeted inhibition of PLK1 has recently been shown to inhibit mutant Kras-dependent tumour growth in both murine and human lung cancer models ^26^. In addition, FOXM1, HELLS, CEP55, and PLK1 are all linked to increased tumour cell aneuploidy ^27,28^, and PLK1 has been shown to induce epithelial-to mesenchymal transition in prostate epithelial cells ^29^. Comparative microarrays were also enriched in genes linked to increased cell cycling (CCNB2, CDKN3, CCNA2) and DNA repair (RAD51, RAD54L, ERCC6L, BRCA1, BRCA2). Activation and disruption of these key signalling pathways are important components of disease progression and metastases in virtually all cancers, and are increasingly exploited by small molecule and biological therpies^30–32^. Separately, UHRF1 is a diagnostic biomarker whose increased expression has been associated with increased lung cancer metastases and cell proliferation^33^, whereas RRM2 is a key regulator of Bcl-2 protein stability, whose overexpression is linked to increased cancer cell resistance to intrinsic apoptosis pathway signalling^34^. DTYMK, which catalyses dTTP biosynthesis, is a necessary component of the recently described Lkb1-null squamous lung tumorigenesis model^35^. Thus, the present study has identified numerous gene expression alterations of likely functional significance in lung cancer ontogeny. The possibility that these represent clinically relevant targets can now be tested using established human lung organoid and air liquid interface models ^36^.

The current study corroborates previous skin, lung, and other epithelial damage models that demonstrate that tissue injury both causes an increased incidence and shortening of the time required for tumour development ^37,38^. In addition, the current results are consistent with recent punctuated equilibrium and macroevolutionary theories of human tumorigenesis (sometimes referred to as the ‘Big bang’ and ‘hopeful monsters’ theories of carcinogenesis, respectively)^39,40^. Although these models differ significantly in estimating how molecular alterations accumulate, each accepts that multiple changes will necessarily occur prior to the establishment of clinically detectable disease (i.e, following tissue injury and repair). In light of these results, the present study supports a model in which gene expression alterations normally associated with airway injury and repair can promote the accumulation of increasingly abnormal lung phenotypes (Figure 6). At some point, a threshold (or rubicon) is crossed that denotes a shift from productive to non-productive repair, culminating in the establishment of early-stage disease. The crossing of this rubicon may simply be due to the severity and/or longevity of airway injury, or may relate to the accumulation of specific molecular alterations such as DNA mutations, chromosomal rearrangements, epigenetic changes, or the inappropriate activation of pathogenic signalling pathways. These observations have significant implications regarding the pathogenesis of human SqCC, and suggest that an increased emphasis on early molecular screening is warranted.

**Figure 6.**
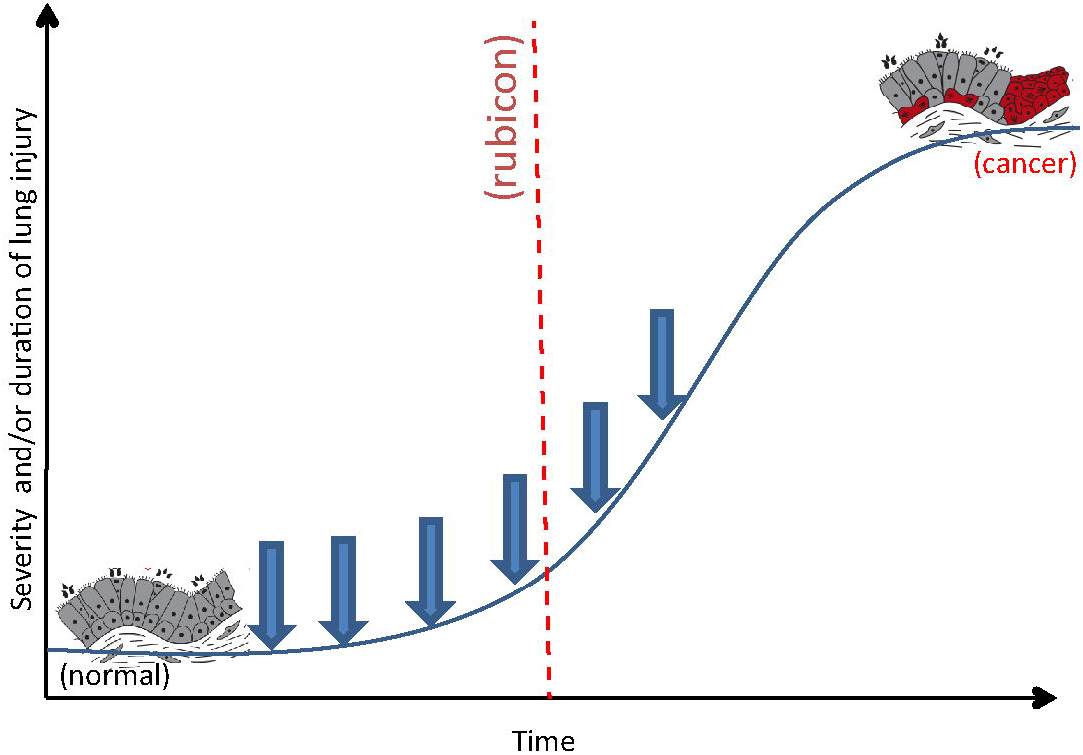
Model linking airway repair and lung cancer. After injury, endogenous airway repair and regeneration triggers expression of conserved signalling pathways and functional biomarkers that promote increased cell proliferation, motility, and reduced epithelial differentiation (blue arrows). Severe and/or chronic lung injury promotes accumulation of additional genetic and epigenetic mutations leading to the establishment of a rubicon, or threshold event that distinguishes productive versus non-productive repair (red line).

Curiously, although several of the genes identified in the present study have previously been identified in established lung cancers, very few have been associated with early stage, preinvasive disease. One reason for this may be the limited number of samples used in previous studies of early human SqCC (n=4, ^6^). In addition, previous experiments have been unable to distinguish molecular alterations that occur due to cigarette smoke exposure, raising the possibility that many gene expression changes were masked due to smoking-induced field-of-injury effects ^10,41^. Indeed, several genes identified in previous studies, but absent in the current dataset analysis, are known to be directly influenced by chronic smoke exposure ^6^. These findings highlight the importance of considering of field-of-injury effects when selecting appropriate controls for investigating early-stage disease.

In summary, I present a collection of gene expression alterations with relevance to human SqCC that were identified using a comparative microarray analysis strategy. It is now possible to assess whether these identified genes may exhibit clinical utility as indicators of SqCC progression and patient mortality. Future studies will also establish the tractability of individual identified genes as novel therapeutic drug targets. While the current study has focused initially on lung SqCC, this comparative microarray approached is easily applicable to additional chronic lung diseases such as fibrosis and COPD. It will also be straightforward to apply this method to the identification of any conserved gene expression changes that are shared across multiple platforms regardless of species, injury models, and disease type.

## Materials and methods

### Gene Microarray Datamining

Gene microarrays for human SqCC (GSE21933, ^42^), murine tracheal polidocanol injury (GSE17268, ^17^) and murine SO2 injury (GSE69056, ^16^) were downloaded from the National Centre for Biotechnology Information (NCBI) Gene Expression Omnibus (GEO), (http://www.ncbi.nlm.nih.gov/geo/). Details of experimental procedures including specifics of murine injury are included in each original research study ^16,17,42^. All GEO datasets included RefSeq Gene ID, gene name, Gene symbol, microarray ID, adjusted P-value, and experimental group logFC relative to corresponding control groups. Experimental groups exhibiting an adjusted p-value of less than 0.05 and a logFC greater than 0.5 (<-0.5 or >0.5) relative to controls following unbiased GEO2R analysis were used in subsequent comparative microarray analyses. For all microarrays, individual samples used to define control and experimental groups are presented in Supplemental Table 1.

### Comparative microarray analysis

To identify shared genes exhibiting differential expression across multiple microarrays a ‘Find Matches’ Visual Basic macro was written for Microsoft Excel which searched for common RefSeq mRNA gene IDs or gene symbols. Details are provided in Supplemental methods. Briefly, gene symbols or gene IDs from two selected datasets were copied into Microsoft Excel, highlighted, and the Visual Basic ‘Find Matches’ macro used to identify genes present in each dataset being compared (defined as shared genes). The logFC of these was then copied into GraphPad Prism, with X and Y values indicating experimental logFC values from each microarray being compared. Genes exhibiting consistent upregulation or consistent downregulation in each microarray being compared were classified as coregulated. To identify genes conserved between airway repair and human SqCC the average logFC of genes selected following comparisons of GSE17268 and GSE69056 (referred to as Average logFC Repair) was compared against the logFC of differentially expressed GSE21933 SqCC genes. To validate the expression of identified genes, I queried the SqCC gene array dataset GSE19188, representing 65 normal and 27 SqCC patient biopsy samples that were independent from GSE21933-derived samples ^19^.

### Kaplan Meier survival analysis

To analyse the effect of individual genes on lung cancer patient mortality I generated survival curves using the online Kaplan-Meier Plotter (Kmplot, http://www.kmplot.com, ^23^). Survival curves comparing individual gene expression versus patient mortality were generated by selecting the indicated Affymetrix gene ID and the ‘Auto select best cutoff’ parameter. Where more than one Affymetrix gene ID was available those IDs with the highest number of associated samples in Kmplot were chosen. Results were derived from the 2015 database release from a total of 1926 (PLK1, POLA2, RABGAP1L) or 1145 patients (WDR78) representing all histology, grade, and stage types. The cutoff value discriminating ‘low’ versus ‘high’ expression, hazard ratio (HR), and logrank P values were generated automatically.

### Tissue preparation, histology, and immunostaining

Human biopsy samples were obtained via fibre-optic bronchoscopy with full informed patient consent and approved by UCLH institutional and national committees, in accordance with relevant guidelines and regulations (UCL Research Ethics committee Approval 06/Q0505/12). Tissues were prepared as previously described ^43^. Briefly, lung samples were formalin fixed overnight at 4 degrees Celsius, paraffin embedded, and sectioned at five microns followed by haematoxylin and eosin (H&E) staining for histological assessment of airway epithelial differentiation. For immunostaining, tissue sections were stained with antibodies directed against the pan-epithelial cytokeratin AE1/3 (DAKO, USA) and PLK1 (Abcam, UK). Sections were dewaxed, blocked using TBS/5%BSA/0.1%T20 for 1h at RT, and incubated overnight at 4C. Secondary antibodies including directly conjugated anti-Rabbit IgG (Alexa 555, Molecular probes, USA) and anti-mouse IgG1 (Alexa 488, Molecular probes) were incubated for three hours at room temperature. Slides were counterstained with 4' 6-diamidino-2-phenylindole (DAPI), washed, and coverslipped using Mowiol 4-88 (Sigma, USA).

### Statistical analysis

The p-value generated in microarray datasets comparing experimental and control groups was generated automatically in Geo2R. Statistical comparisons of gene expression were performed using GraphPad Prism Software (La Jolla, CA). An unpaired T test with Mann-Whitney correction was used to compare individual genes in control vs 2d polidocanol, control vs SO2 injury, and normal versus SqCC samples. An exact statistical test to establish the significance of intersections amongst independent lists of genes was established according to the methods outlined in Natarajan et al and is presented in supplemental data ^18^.

## Acknowledgements

I gratefully acknowledge Dr Jeremy George, Prof Sam Janes, and Ms Bernadette Carroll for providing bronchoscopic tissue biopsy samples used for immunostaining. I thank Ms Pooja Seedhar for biopsy sample embedding and preparation, and Drs Vitor Teixeira and Adam Pennycuik for helpful discussions regarding microarray analysis and interpretation. I acknowledge the support of Dr Andy Blanchard for a discussion regarding the clinical relevance of this study, and Dr Trevor Graham for a discussion regarding the interpretation of his recent ‘Big Bang’ theory of carcinogenesis.

## Additional Information, Competing Financial Interests

There are no competing financial interests related to this manuscript.

## Author contributions

AG designed, performed, wrote, and reviewed all components of this manuscript.

